# A comparison of DNA stains and staining methods for Agarose Gel Electrophoresis

**DOI:** 10.1101/568253

**Authors:** Andie C. Hall

**Affiliations:** Core Research Laboratories. Natural History Museum, London SW75BD.

**Keywords:** Agarose gel electrophoresis, DNA stain, fluorescent dye

## Abstract

Nucleic acid stains are necessary for Agarose Gel Electrophoresis (AGE). The commonly used but mutagenic Ethidium Bromide is being usurped by a range of safer but more expensive alternatives. These safe stains vary in cost, sensitivity and the impedance of DNA as it migrates through the gel. Modified protocols developed to reduce cost increase this variability. In this study, five Gel stains (GelRed™, GelGreen™, SYBR™ safe, SafeView and EZ-Vision®In-Gel Solution) two premixed loading dyes (SafeWhite, EZ-Vision®One) and four methods (pre-loading at 100x, pre-loading at 10x, precasting and post-staining) are evaluated for sensitivity and effect on DNA migration. GelRed™ was found to be the most sensitive while the EZ-Vision® dyes and SafeWhite had no discernible effect on DNA migration. Homemade loading dyes were as effective as readymade ones at less than 4% of the price. This method used less than 1% of the dye needed for the manufacturer recommended protocols. Thus, with careful consideration of stain and method, Gel stain expenditure can be reduced by over 99%.

## 1. INTRODUCTION

Nucleic acid stains are intercalating dyes which bind to DNA and fluoresce under UV light. Stains can be added to loading dye and then mixed with the DNA prior to running a gel (pre-loading), added to the gel itself so the DNA picks up the stain as it migrates (precasting) or the gel can be soaked in a staining solution after AGE has finished (post-staining).

Intercalating dyes change the charge and flexibility of DNA molecules and add weight, altering movement through the gel (Sigmon and Larcom 1996; Miller et al. 1999; Huang and Fu 2005; Huang et al. 2010).The post-staining method is therefore the most accurate way to size DNA fragments but it is time consuming and costly as more stain is needed.

Preloading is not recommended by dye manufacturers but uses considerably less stain than standard protocols, making it a much cheaper method. The response from manufacturers is new, commercially available, ready-made DNA loading dyes, to replace the “homemade” versions. These dyes are considerably more expensive per sample than the homemade equivalents.

Precasting with Ethidium Bromide (EtBr) is common but due to its mutagenicity it has been phased out of many labs. Alternatives vary greatly in price and sensitivity although all claim equal or greater sensitivity than EtBr. Most are designed for use in the same way as EtBr (Precast or post-stain).

Biotium state that GelRed™ and GelGreen™ have a much greater mass than EtBr so that they cannot cross cell membranes. This makes them nontoxic and nonmutagenic but also slows the DNA as it moves through the gel (Couto et al 2013). Biotium’s website states “because GelRed™ and GelGreen™ are high affinity dyes designed to be larger dyes to improve their safety, they can affect the migration of DNA” (Biotium 2018). Crisafuli et al (2015) suggest GelRed™ to be a bis-intercalator, formed from two cross-linked Ethidium Bromide molecules. If this is so, they state GelRed should be twice as sensitive as EthBr since “the GelRed assay will have approximately twice as many DNA-bound sites”. They found a marked effect on DNA contour length as well as weight, which would affect the DNA’s movement through the gel. Nath et al (2000) describe a similar effect for SYBR™GreenI at high concentrations. Despite this, Huang et al (2010) and Bi et al (2011) reported no effect on DNA mobility with GelRed at 100x concentration in a loading dye.

GelGreen™ can be visualised with UV or blue light (such as on a dark reader), for easier gel excision but in all other practical respects, is the same as GelRed™.

AMRESCO claim that EZ-Vision® has similar sensitivity to EtBr, but without any effect on electrophoretic mobility. A number of ready-made loading dye versions are available, which are added to each sample before electrophoresis. Tested here are EZ-Vision®In-Gel Solution (designed for precasting) and EZ-Vision®One (a pre-loading dye; added to each sample before electrophoresis).

SafeView from NBS bio and SYBR™safe from Invitrogen are claimed to be as sensitive as EtBr but no information on DNA migration could be found. Both may be visualised with either UV or blue light. SafeWhite is a pre-loading version of SafeView.

Published comparative studies of DNA stains compare relatively old versions of stains, do not compare staining methods, or include little reference to band-sizing problems. No independent reviews of premixed loading dyes, or comparisons with homemade versions were found, presumably as they are relatively new products. In this study, 5 Gel stains (GelRed™, GelGreen™, SafeView, SYBR® safe, and EZ-Vision®In-Gel Solution) and 2 readymade loading dyes (SafeWhite, and EZ-Vision® One) are evaluated for sensitivity and migration using pre-loading at 2 concentrations, precasting and post-staining methods.

## 2. MATERIALS AND METHODS

In all cases, a 1% agarose 1xTAE gel was used, run at 90volts for one hour.

### 2.1 Pre-loading method

GelRed™10,000x in DMSO, GelGreen™10,000x in DMSO, EZ-Vision®In-Gel Solution10,000x in DMSO, SafeView 10,000x and SYBR™safe 10,000x in DMSO were added to blue loading dye (0.25% Bromophenol blue, 0.25% Xylene cyanol, 30% glycerol solution), and to the loading dye supplied with Lambda DNA/HindIII marker™ (ThermoScientific™) at a 1:500 and 1:50 dilution. 1μl of loading dye was then added to 1μl of each PCR product to give final stain concentrations of 10x and 100x. For the Lambda marker, 0.6μl of loading dye was added per lane, as per manufacturer’s instructions. Stain was also added to the markers Hyperladder 1kb™ and Hyperladder 4™ (Bioline) to give final concentrations of 10x and 100x.

0.2μl of EZ-Vision®One was mixed with 1μl each PCR product, and 0.6μl to the lambda marker before loading. It was mixed with Hyperladder 1kb™ and Hyperladder 4™ to a 1:5 dilution before loading, as per manufacturer’s instructions.

2μl of SafeWhite was mixed with 1μl each PCR product, and to the Lambda DNA/HindIII marker™ before loading. It was mixed with Hyperladder 1kb™ and Hyperladder 4™ to a 1:5 dilution before loading, as per manufacturer’s instructions.

### 2.2 Precast

All stains were used as follows: 5μl of stain was added to 50ml of 1% molten agarose in TAE before casting (10,000x dilution), as per manufacturer’s instructions.

### 2.3 Post-Stain

Post-staining was carried out in accordance with manufacturers’ instructions, specifically; GelRed™ and GelGreen™: 15μl of stain was added to 50ml water. The gel was submerged in the stain solution, wrapped in aluminium foil and placed on an orbital shaker for 30 minutes.

EZ-Vision®In-Gel Solution: 12.5μl of stain was added to 50ml of 100mM NaCl. The gel was submerged in the stain solution, wrapped in aluminium foil and placed on an orbital shaker for 30 minutes. The stain solution was replaced with water for 2× 10mins on the shaker.

SafeView: 12.5μl of stain was added to 50ml of 1×TAE. The gel was submerged in the stain solution, wrapped in aluminium foil and placed on an orbital shaker for 20 minutes.

SYBR™safe: 5μl of stain was added to 50ml of 1×TAE. The gel was submerged in the stain solution, wrapped in aluminium foil and placed on an orbital shaker for 30 minutes.

Gels were photographed using an Alpha Imager HP (Alpha Innotech), on a UV transilluminator 365nm with either an Ethidium Bromide or SYBR™ filter (whichever produced a clearer picture) and on an Invitrogen SafeImager™ with the supplied amber filter. Exposure times were adjusted for each gel to give the clearest image. 10x Preloading gels were imaged with all 3 methods to ascertain the best imaging method for subsequent steps.

## 3. RESULTS

The preloading method used significantly less stain than the standard precasting and post-staining methods so was much cheaper (Table 1). But, preloading produced few to no visible bands for SafeView and SYBRsafe (table 1. Fig. 1, 2), whilst almost all bands were visible when preloading with the Biotium stains (figs 3, 4. table 1). SafeView was the least sensitive stain when precasting and bands in the post-stained gel were less bright and clear than the other stains (except SYBRSafe) (fig.1). Gels precast with SYBRsafe were clear and smaller fragments could be accurately sized, however there were size discrepancies for bands over 4kb (Table 1. Fig.2). The post-stained gel suffered from speckling.

**Fig. 1.**
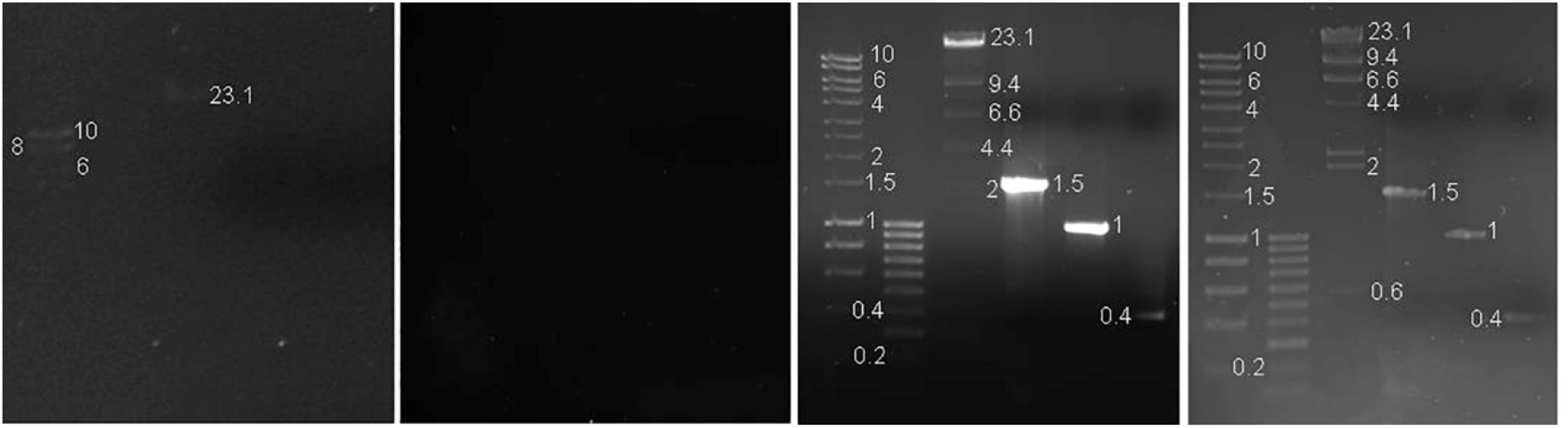
SafeView Left: Preload 10 x Left Middle: Preload 100x Right Middle: Precast Right: Post-stain. Photographed on a UV transilluminator 365nm with an EtBr filter. Lane 1: 1μl of Hyperladder 1kb™ (Bioline) 2: 1μl of Hyperladder IV ™ (Bioline) 3: 0.3μg Lambda DNA HindIII marker2 ™ (ThermoScientific™) 4: 580ng of PCR product 5: 27ng of PCR product 6: 8ng of PCR product (as measured on a Qubit ™ Fluorometer)

**Fig. 2.**
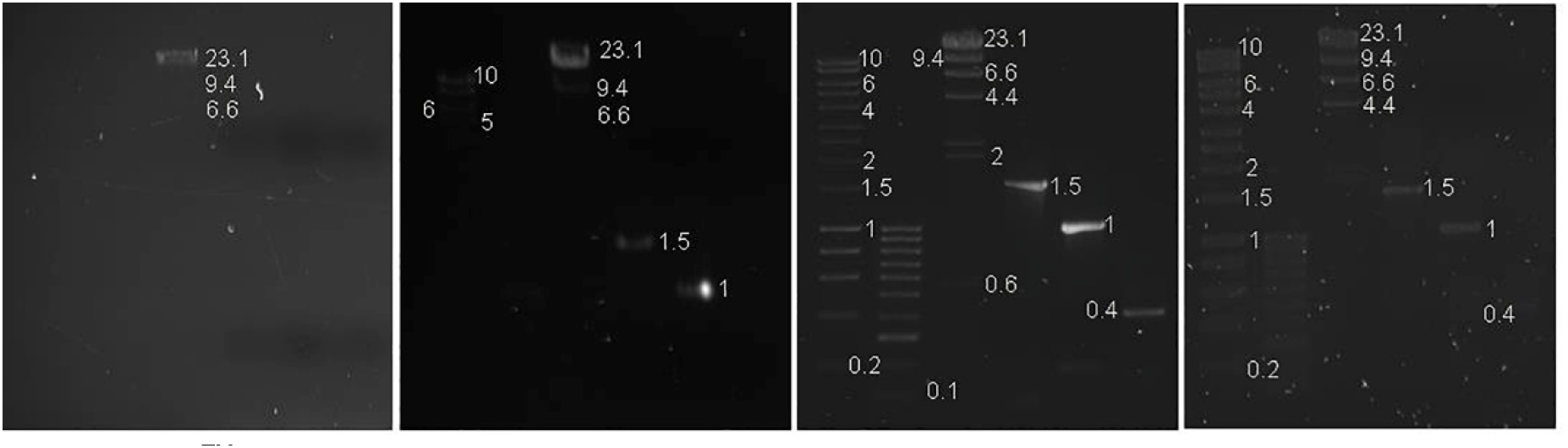
SYBR™Safe Left: Preload 10 x Left Middle: Preload 100x Right Middle: Precast Right: Post-stain.Photographed on a UV transilluminator 365nm with a SYBR™ filter. Lane 1: 1μl of Hyperladder 1kb™ (Bioline) 2: 1μl of Hyperladder IV ™ (Bioline) 3: 0.3μg Lambda DNA HindIII marker2 ™ (ThermoScientific™) 4: 580ng of PCR product 5: 27ng of PCR product 6: 8ng of PCR product (as measured on a Qubit ™ Fluorometer)

**Fig. 3.**
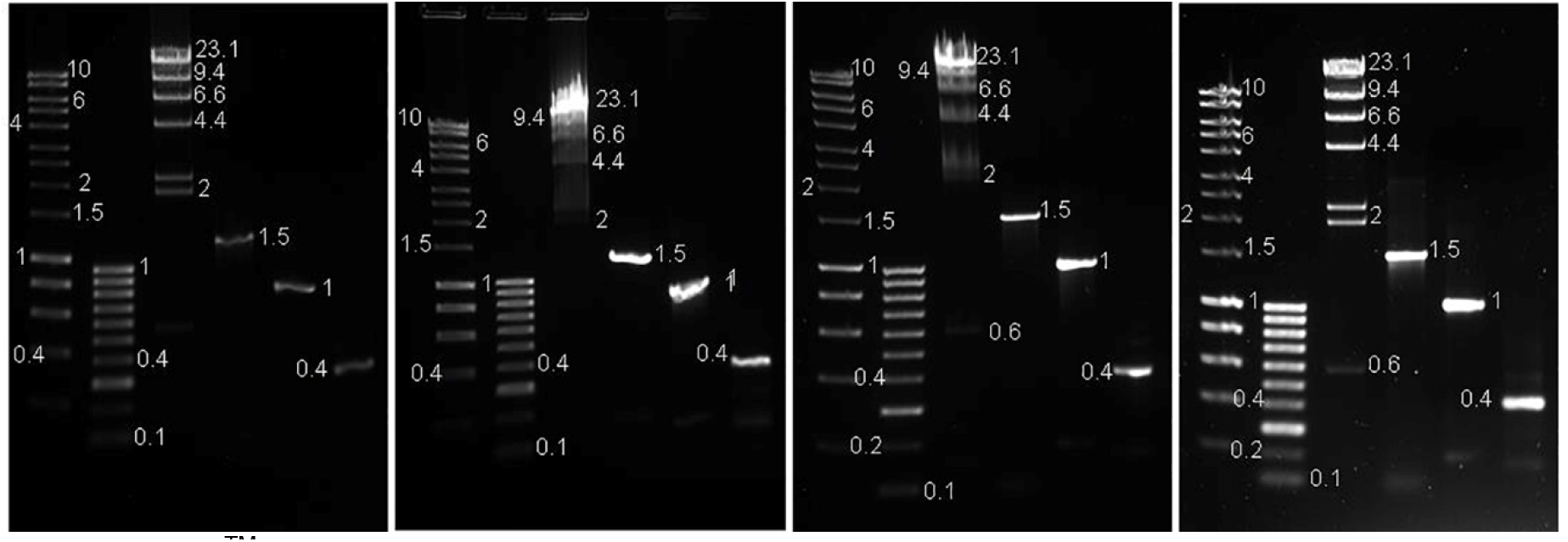
GelRed™ Left: Preload 10x. Left Middle: Preload 100x. Right Middle: Precast. Right: Post-stain. Photographed on a UV transilluminator 365nm with an EtBr filter. Lane 1: 1μl of Hyperladder 1kb™ (Bioline) 2: 1μl of Hyperladder IV ™ (Bioline) 3: 0.3μg Lambda DNA HindIII marker2 ™ (ThermoScientific™) 4: 580ng of PCR product 5: 27ng of PCR product 6: 8ng of PCR product (as measured on a Qubit ™ Fluorometer)

**Fig. 4.**
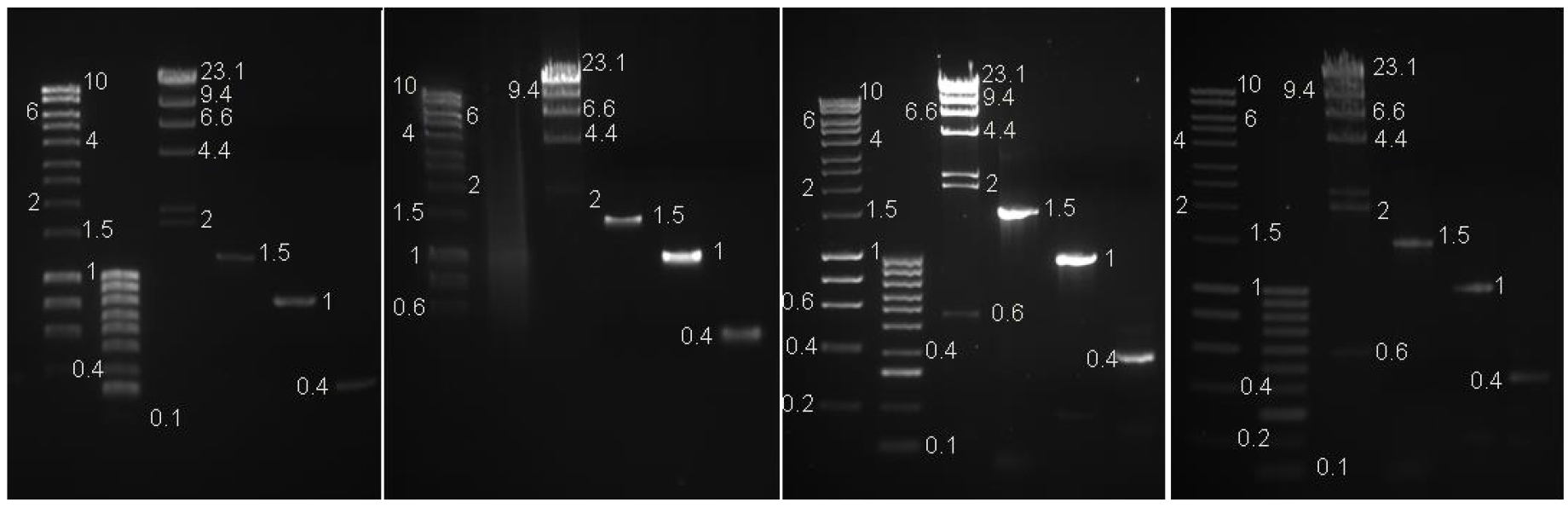
GelGreen™ Left: Preload 10x Left Middle: Preload 100x Right Middle: Precast Right: Post-stain. Photographed on an Invitrogen SafeImager ™ with an amber filter. Lane 1: 1μl of Hyperladder 1kb™ (Bioline) 2: 1μl of Hyperladder IV ™ (Bioline) 3: 0.3μg Lambda DNA HindIII marker2 ™ (ThermoScientific™) 4: 580ng of PCR product 5: 27ng of PCR product 6: 8ng of PCR product (as measured on a Qubit ™ Fluorometer)

**Table 1.** Results summary. Costs are for comparison only and based on volume of stain used in this study for six lanes on one 50ml gel, using list prices in Pound Sterling (GBP) on 17.12.19 except the EZVision stains which are no longer available in the UK- *Prices for EZ vision are in USD

| Stain | Method | Visible bands | Consistent DNA migration | Total cost of stain per assay (pence*) |
| --- | --- | --- | --- | --- |
| EZ vision in gel solution | 10x Preloading | 30/35 | Yes | 0.2 |
|  | 100x preloading | 30/35 | Yes | 2.0 |
|  | Precasting | 34/35 | Yes | 109.3 |
|  | Post staining | 33/35 | Yes | 273.3 |
| GelGreen | 10x Preloading | 30/35 | No for $\geq 0.4\text{kb}$ | 0.2 |
|  | 100x preloading | 21/35 | No for 1.5-2.5kb | 1.9 |
| | Precasting | 34/35 | No for $\geq 0.4\text{kb}$ | 105.0 |
|  | Post staining | 34/35 | Yes | 315.0 |
| GelRed | 10x Preloading | 34/35 | No for $\geq 0.4\text{kb}$ | 0.2 |
|  | 100x preloading | 33/35 | No | 2.2 |
| | Precasting | 34/35 | No for $\geq 0.4\text{kb}$ | 121.0 |
|  | Post staining | 34/35 | Yes | 363.0 |
| SafeView | 10x Preloading | 4/35 | NA | 0.04 |
|  | 100x preloading | 0/35 | NA | 0.4 |
| | Precasting | 32/35 | No for $\geq 1.5\text{kb}$ | 21.0 |
|  | Post staining | 33/35 | Yes | 52.5 |
| SYBR safe | 10x Preloading | 9/35 | Yes | 0.1 |
|  | 100x preloading | 3/35 | NA | 1.4 |
| | Precasting | 34/35 | No for $\geq 4\text{kb}$ | 76.9 |
|  | Post staining | 30/35 | Yes | 76.9 |
| EZ vision One | Preloading | 30/35 | Yes | 4.0 |
| SafeWhite | Preloading | 31/35 | Yes | 18.5 |

The Biotium dyes showed the greatest sensitivity for all the methods with exception of preloading GelGreen™ at 100x but had a marked effect on DNA migration-the only way to accurately size fragments with these stains is post-staining. Larger bands stained with 100x Biotium preloading dyes also showed smearing (Table 1, Figs 3,4). Hyperladder IV™ became a long smear without discernible bands in the 100x GelGreen™ preloading treatment (Fig 4). The solution itself became stringy and separated into layers. This test was repeated with fresh ladder and dye, but with the same result. The 100x loading dyes showed reduced sensitivity compared to 10x preloading for SYBR™Safe and SafeView but had no effect on EZ-Vision®In-Gel Solution (table 1).

Although the EZ-vision® dyes were slightly less sensitive than GelRed™ and GelGreen™; neither affected DNA migration. The same bands were visible with the EZ-Vision®In-Gel Solution preloading methods as with EZ-Vision®One which is considerably more expensive (Table 1, Fig 5, 6). SafeWhite was slightly more sensitive than the EZVision dyes and also showed no discernible effect on DNA migration (Fig 6).

**Fig. 5.**
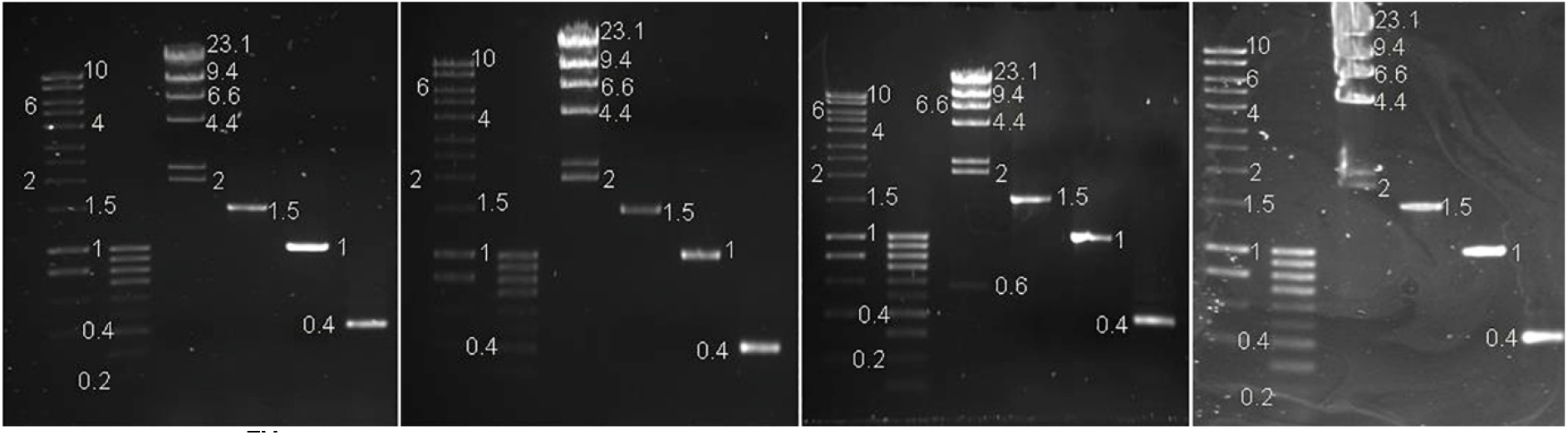
EZ-vision™In-Gel solution. Left: Preload 10x Left Middle: Preload 100x Right Middle: Precast Right: Post-stain.Photographed on a UV transilluminator 365nm with a SYBR™ filter. Lane 1: 1μl of Hyperladder 1kb ™ (Bioline) 2: 1μl of Hyperladder IV ™ (Bioline) 3: 0.3μg Lambda DNA HindIII marker2 ™ (ThermoScientific™) 4: 580ng of PCR product 5: 27ng of PCR product 6: 8ng of PCR product (as measured on a Qubit ™ Fluorometer)

**Fig. 6.**
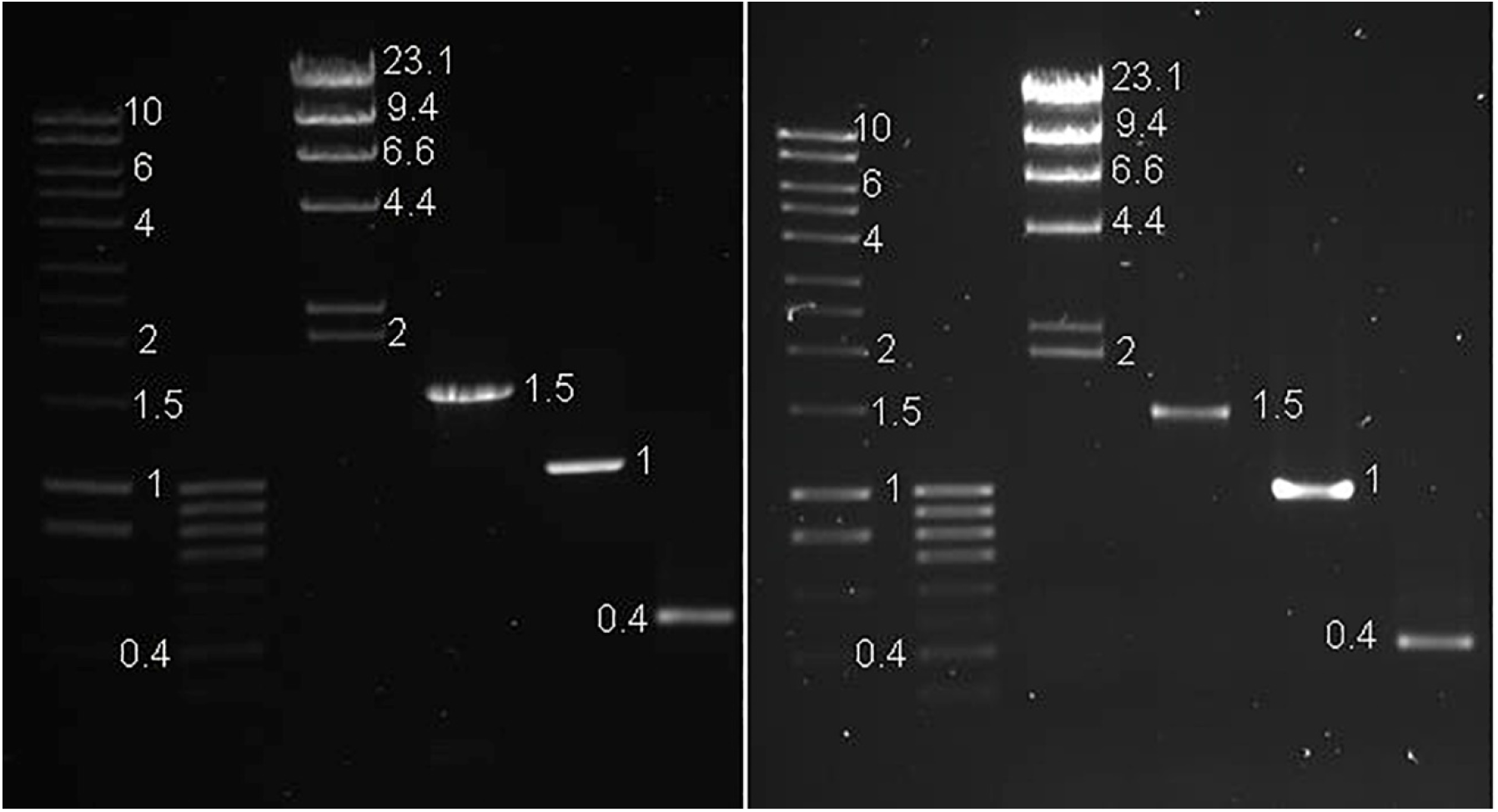
Left: EZ-Vision®One Right: SafeWhite. Photographed on a UV transilluminator 365nm with a SYBR™ filter. Lane 1: 1μl of Hyperladder 1kb™ (Bioline) 2: 1μl of Hyperladder IV™ (Bioline) 3: 0.3μg Lambda DNA HindIII marker2 (ThermoScientific™) 4: 580ng of PCR product 5: 27ng of PCR product 6: 8ng of PCR product (as measured on a Qubit ™ Fluorometer)

GelGreen™ was the only stain to work under blue light (Fig 7).

**Fig. 7.**
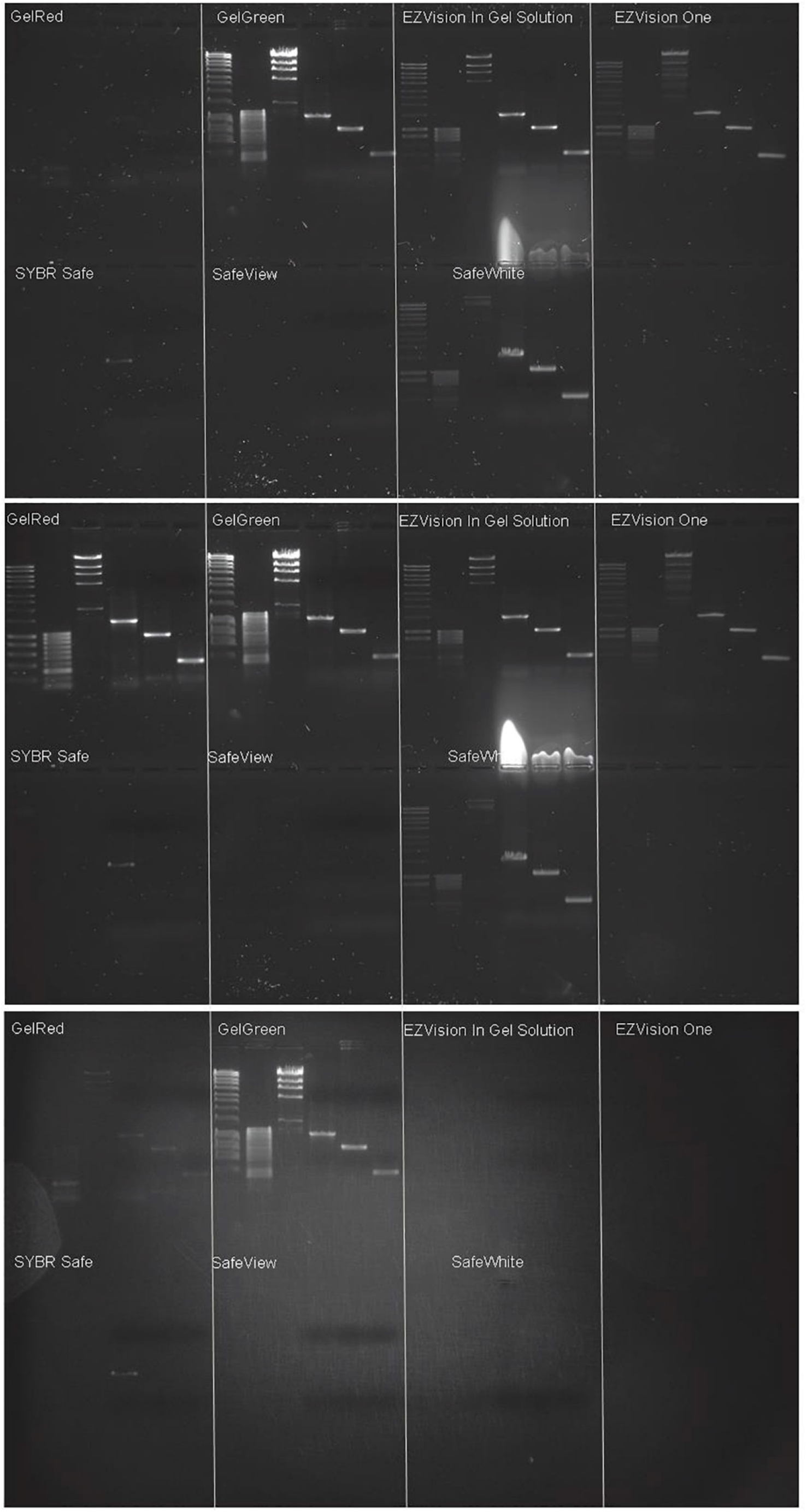
Preload 10x. Top panel: UV transilluminator 365nm with a SYBR™ filter. Centre panel: UV transilluminator 365nm with an EtBr filter. Bottom panel: Invitrogen SafeImager with amber filter. Exposure times vary. Lane 1: 1μl of Hyperladder 1kb™ (Bioline) 2: 1μl of Hyperladder IVTM (Bioline) 3: 0.3μg Lambda DNA HindIII marker2 (ThermoScientific™) 4: 580ng of PCR product 5: 27ng of PCR product 6: 8ng of PCR product

## 4. DISCUSSION

### 4.1 Method comparison

Adding DNA stains to loading dyes before AGE significantly reduces the amount of stain used, and thus represents a huge cost saving (Table 1) compared to the standard precasting and post-staining methods. In our example, making a 10x preloading dye used less than 1% of the stain used for precasting, however, the method simply didn’t work for SafeView and SYBRsafe and the Biotium stains affected DNA migration. Mobility was also affected by Biotium stains when using the precast protocol though to a lesser extent (Table 1, Figs 3,4). There was disagreement between bands of the same size when using GelRed™ in both preloading dyes, disagreeing with the conclusions of Huang et al (2010) and Bi et al (2011). Larger bands stained with 100x Biotium preloading dyes also showed smearing, possibly due to overloading (Fig 3, 4. Biotium 2019). The higher concentration of GelGreen™ in the loading dye had an unexpected effect on one of the ladders; Hyperladder IV™ appeared to degrade, separating into layers and becoming stringy-resulting in a long smear without discernible bands (Fig 4). Biotium were contacted but couldn’t offer an explanation (Biotium Techsupport, pers.comm). Whilst Biotium now offer a preloading version of Gelred, no such product exists for GelGreen. Increasing the stain to 100x was also detrimental to SYBR™Safe and SafeView, resulting in reduced sensitivity (table 1).

In most cases, post-staining was the most sensitive and accurate method for DNA band sizing (table 1). It is also the most expensive due to the volume of stain used, though manufacturers state that staining solution can be reused 3 times to reduce costs. It is also considerably more time consuming than the others, with soaking and wash steps adding 20-50mins to the protocol.

Whilst precasting seems to represent a halfway-house in terms of cost, time, sensitivity and ability to size DNA accurately; all factors vary with the stain used, so each should be considered individually when choosing a staining method.

### 4.2. Stain comparison

The Biotium dyes were the most sensitive but the only way to accurately size fragments with these stains is post-staining (Fig 3, 4 Table 1).

Preloading with EZ vision dyes was slightly less sensitive than GelRed™ and GelGreen™ but the other methods were comparable and DNA migration appeared unaffected.

SYBR™safe did not work as a preloading dye but precast gels were clear and smaller fragments could be accurately sized, however there were size discrepancies for bands over 4kb (Table 1. Fig.2). The post-stained gel suffered from speckling so was difficult to read. “Many whitening agents used in clothing, as well as some fungi and bacteria, fluoresce at the same wavelengths as SYBR™Safe DNA gel stain. These contaminants, within or on the surface of the gel, may produce speckling” (ThermoFisher Scientific 2018).

Pre-loading with SafeView produced few to no visible bands (Fig. 1 & 5). Migration of the stain in the precast gels caused the top to appear washed-out while the bottom was too dark to see the smallest bands. Whilst this can occur with all stains, all gels in this paper were run in the same way and stain migration for the other stains was not noticeable. Bands in the post-stained gel were less bright and clear than the other stains (except SYBRSafe) but as the cheapest stain tested, increasing the volume of stain per gel may be still be economical. NBS Biologicals no longer recommend SafeView for post-staining, suggesting users purchase *SafeView Plus* instead (NBS biologicals 2018).

The commercially made pre-loading dyes EZ-Vision®One and SafeWhite were extremely easy to use and had no effect on DNA migration (Fig 7). However, the 10x preloading dyes made with GelRed™, GelGreen™ and EZ-Vision®In-Gel Solution were more sensitive and considerably cheaper (Table 1). Since bands stained with the EZ-Vision®In Gel solution preloading method also ran true to size, the only benefit of purchasing a readymade loading dye seems to be convenience.

Whilst GelGreen™ was best visualized with blue light, bands were as bright and clear with a UV light making it the only stain to work under both lights (Fig 7).

### 4.3 Concluding remarks

Whilst not recommended by stain manufacturers, the 10x preloading method can reduce stain expenditure by over 99% with little to no loss in sensitivity. Increasing the stain concentration to 100x was detrimental as well as more expensive. DNA band-sizing problems can be mitigated with careful consideration of method and brand, with no need to purchase readymade preloading dyes, other than convenience. Based on cost, sensitivity and stability, our lab routinely uses our own recipe preloading dye made with a 10x concentration of GelRed™. However this method with EZ-Vision®In-Gel Solution is preferred when accurate sizing of fragments is necessary. When both high sensitivity and accurate sizing are desired, post-staining with a Biotium dye is recommended, reusing the stain solution where possible to reduce costs.

## ACKNOWLEDGEMENTS

This study was funded by the Core Research Laboratories, Natural History Museum, London.

## Notes

#### Summary of Updates

Results and discussion separated. Prices updated. Figure 4 amended (original version images for 10x loading dye and 100x loading dye were the wrong way round)

